# Did you see the sound? A Bayesian Perspective on Crossmodal Perception in Low Vision

**DOI:** 10.64898/2025.12.24.696433

**Authors:** Ailene Y. C. Chan, Noelle R. B. Stiles, Carmel A. Levitan, Armand R. Tanguay, Shinsuke Shimojo

## Abstract

Multisensory integration is often assumed to increase when visual input is degraded, yet it remains unclear whether low vision enhances susceptibility to cross-modal illusions or whether such effects depend on local variations in visual reliability. We tested low vision and sighted control participants on the Double Flash Illusion across 24 visual-field locations while separately measuring flash-detection accuracy. Both groups showed the expected auditory-driven increase in perceived flash numerosity, but only sighted controls reliably experienced the classic “double-flash” percept. Illusion strength did not vary with eccentricity; instead, it was strongly predicted by local flash detection accuracy, indicating that sound-induced percepts depend on the availability of a reliable visual signal. Bayesian Causal Inference modeling revealed substantially weaker and more variable fits for low vision observers, with poorer fits associated with reduced visual sensitivity. Although model parameters did not differ significantly between groups, the similarity in estimated visual noise likely reflects model limitations rather than true equivalence in sensory precision. Together, these findings show that low vision does not globally amplify audiovisual interactions; rather, auditory enhancement depends on local visual reliability, and degraded vision leads to weaker alignment with Bayesian-optimal predictions.

## 1 Introduction

In every waking moment, the nervous system must translate incomplete and noisy sensory input into a coherent percept. When multiple senses are simultaneously perceived the brain must also infer whether multisensory information originates from a single cause or from separate events. A healthy nervous system accomplishes this by weighting sensory cues according to their reliability and by incorporating prior beliefs (or heuristics) about how sensory events unfold [Ernst and Banks, 2002, Körding et al., 2007]. In this framework, perception reflects the statistically optimal combination of cues given their uncertainties and the observer’s expectations.

But what happens when one of these sensory channels is degraded? While multisensory integration has been well characterized in healthy observers, much less is known about how these computational principles operate when visual input is compromised. Low vision offers a particularly informative case: rather than a complete loss of vision, patients retain islands of intact visual function interspersed with regions of reduced sensitivity. This raises an important question: Is multisensory integration globally altered in low vision, or does it adjust locally according to the reliability of the visual signal at each region of the visual field?

Neural evidence suggests that the visually impaired brain undergoes cross-modal reorganization, with auditory and tactile signals recruiting occipital cortex even when some visual function remains [Cunningham et al., 2015, Norman and Thaler, 2019]. Anatomical studies further reveal direct projections from auditory cortex to early visual regions [Beer et al., 2011, Falchier et al., 2002, Quinones et al., 2022], especially representing peripheral visual space. These findings raise the possibility that cross-modal influences on perception may be enhanced in low vision. Yet, whether such enhancement depends on global differences in sensory processing (e.g., reduced weighting of vision) or local differences driven by the visibility of specific retinal regions remains unresolved.

To address this gap, we used the Double Flash Illusion [Shams et al., 2000], a classic demonstration in which auditory beeps alter the perceived number of visual flashes. By presenting the illusion at 24 locations across the visual field in both low vision and sighted control participants, we aimed to answer three questions: (i) Are low vision participants more susceptible to sound-induced enhancements in perceived flash count compared to sighted controls? (ii) Do multisensory interactions vary across the visual field, particularly in peripheral regions? (iii) Most importantly, does illusion strength depend on local visual sensitivity—that is, does the illusion become stronger at locations where visual detection is reduced?

These questions allow us to disentangle whether multisensory processing is globally altered in low vision or whether it flexibly adapts to local variations in visual reliability across the retina. By combining behavioral measures with Bayesian Causal Inference modeling, we further examined whether any group differences arise from changes in sensory noise, prior expectations, or causal inference strategies.

## 2 Methods

### 2.1 Human Participants

We first recruited 21 neurotypical participants (11 females; mean age = 26.3 years, range = 18–40 years) from the California Institute of Technology community. All had normal or corrected-to-normal vision (20/20) and hearing. Recruitment took place during the early phase of the study, when COVID-19 restrictions limited access to external participants.

We later recruited 25 individuals with low vision after obtaining written informed consent, either through an e-reader or verbally through the experimenter. Four participants did not complete the full experimental protocol and were excluded from data analysis. The final low-vision cohort included 21 participants (8 females; mean age = 51.5 years, range = 25–81 years), all with normal or corrected-to-normal hearing. Demographic and clinical details are shown in Table 1.

**Table 1:** Demographic and Clinical Information of Low Vision Cohort.

| Participant | Diagnosis | Age<br>(Years) | Sex | Age of Onset<br>(Years) | Duration<br>(Years) |
| --- | --- | --- | --- | --- | --- |
| LV001 | Retinitis pigmentosa | 65 | F | 31 | 34 |
| LV002 | Microphthalmia | 43 | M | 0 | 43 |
| LV003 | NAION, right eye | 74 | M | 68 | 6 |
| LV005 | Congenital cataracts | 69 | M | 0 | 43 |
| LV007 | Macular degeneration | 70 | F | 68 | 2 |
| LV008 | Choroideremia | 33 | M | 23 | 10 |
| LV009 | Stargardt's macular<br>degeneration | 28 | M | 14 | 14 |
| LV011 | Left Phthisis Bulbi,<br>Right cystoid macular<br>edema, uveitis | 66 | F | 24 | 42 |
| LV012 | Glaucoma, right eye<br>total blind | 56 | M | 42 | 14 |
| LV013 | Glaucoma, right eye<br>total blind | 40 | M | 35 | 5 |
| LV015 | Retinitis pigmentosa,<br>detached retina, right<br>eye only | 57 | M | 11 | 46 |
| LV016 | Chronic Relapsing<br>Inflammatory Optic<br>Neuritis (CRION) | 52 | F | 36 | 16 |
| LV017 | Enlarged optic nerves | 33 | M | 27 | 6 |
| LV018 | Dominant optic atro-<br>phy | 25 | F | 12 | 13 |
| LV019 | Glaucoma, left eye<br>only | 81 | M | 76 | 5 |
| LV020 | Retinal degeneration | 42 | M | 0 | 42 |
| LV021 | Retinitis Pigmentosa | 41 | F | 21 | 20 |
| LV022 | NAION | 64 | M | 56 | 8 |
| LV023 | Undeveloped optic<br>nerve | 57 | F | 0 | 57 |
| LV024 | Optic nerve damage<br>due to trauma | 42 | F | 3 | 39 |
| LV025 | Microphthalmia | 44 | M | 0 | 44 |

After testing the low-vision group, we recruited an additional cohort of age-matched neurotypical controls to better match the age distribution of the low-vision participants. Five participants from the original control cohort were retained because their ages already fell within the target range of the low-vision group and therefore met the age-matching criteria. To complete the age-matched sample, 16 new control participants were recruited, resulting in a final control cohort of 21 participants (15 females; mean age = 51.0 years, range = 24–80 years).

Participants were compensated at $25/hour. All procedures were approved by the institutional review board at the California Institute of Technology (IRB IR20-1025).

### 2.2 Simuli and conditions

#### 2.2.1 Visual Flash Detection Task

Visual stimuli were circular white discs (2*^◦^* in diameter) presented on a black background at 100% contrast for one frame (16.7 ms at 60 Hz). Auditory stimuli were pure tones (800 Hz, 7 ms duration). Stimuli appeared at 24 possible locations, corresponding to eight equally spaced positions along the circumference at 5*^◦^*, 10*^◦^*, and 15*^◦^* eccentricities from central fixation (Fig. 1A). Two audiovisual conditions were tested in this task, following the naming convention FXBY, where X denotes the number of flashes and Y denotes the number of beeps such as F0B0 (no flashes or beeps) and F1B0 (one flash, no beeps). The stimulus timing for these conditions is illustrated in Fig. 1B–C.

**Figure 1:**
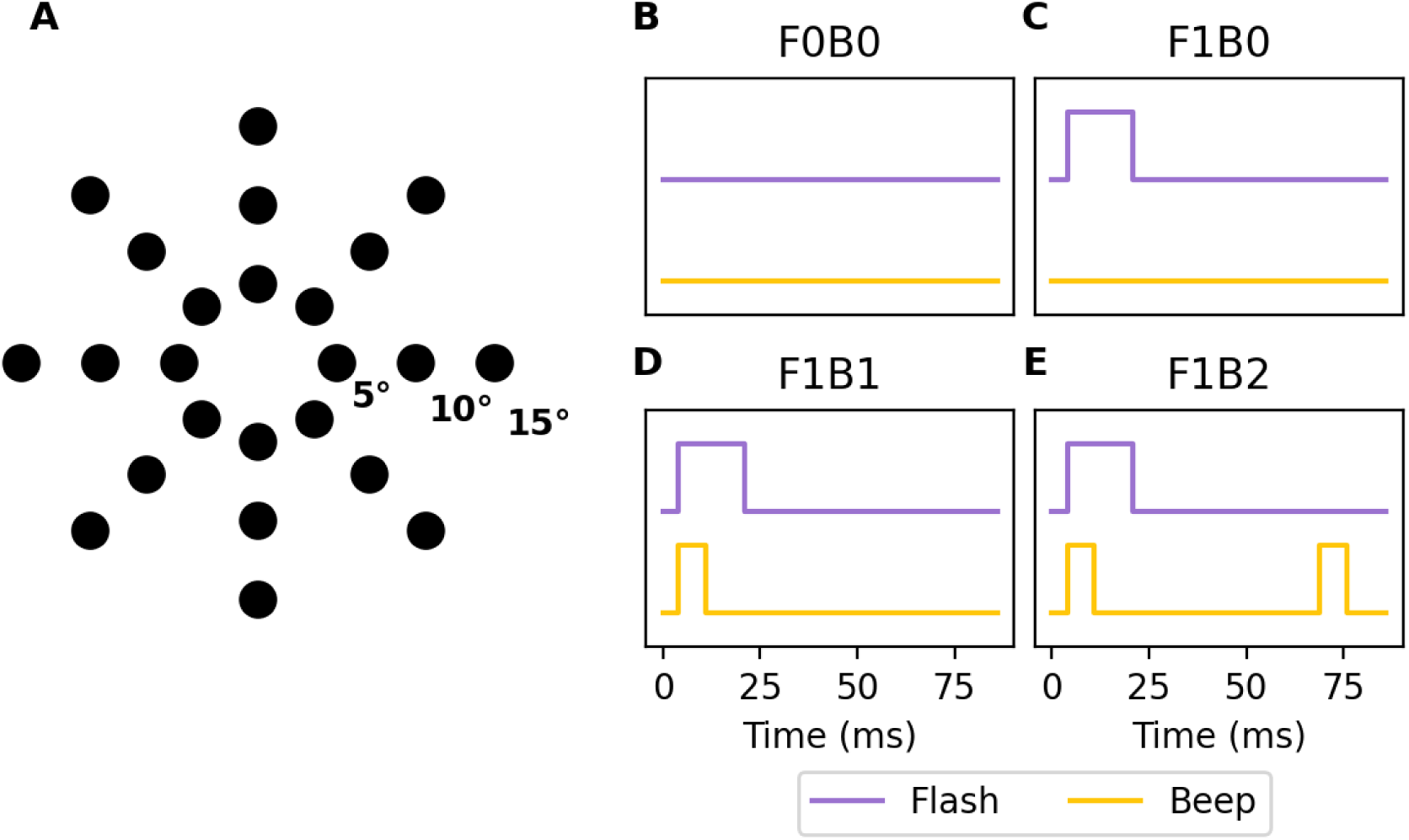
Stimuli and conditions. **(A)** Stimulus locations corresponding to eight equally spaced positions along the circumference at 5*^◦^*, 10*^◦^*, and 15*^◦^* eccentricities from central fixation. **(B-E)** Schematic of the four main experimental conditions, named using an FXBY convention (*X* = number of flashes, *Y* = number of beeps). When paired, visual flashes (16.7 ms) and auditory beeps (7 ms) had simultaneous onsets.

#### 2.2.2 Double Flash Task

This task used the same visual and auditory stimuli and stimulus locations as described above (Fig. 1A). Two audiovisual conditions were tested: F1B1 and F1B2. The F1B2 condition, also known as the Double Flash condition, elicits the perception of a second illusory flash induced by the second auditory beep. Timing of the stimuli is shown in Fig. 1D–E.

#### 2.2.3 Beep Detection Task

This task also used the same visual and auditory stimuli and stimulus locations as the Visual Flash Detection Task (Fig. 1A). Six audiovisual combinations of flashes (0 or 1) and beeps (0, 1, or 2) were tested. Stimulus timing followed the same parameters as described above (see Fig. 1B–E).

### 2.3 Experimental Design

#### 2.3.1 Visual Flash Detection Task

The task was performed monocularly for each eye in randomized order, with the non-tested eye covered by an eyepatch. Participants indicated whether they detected a flash (*Yes*: press left arrow button; *No*: press right arrow button). The F1B0 condition was presented five times per stimulus location, and the F0B0 condition served as a single catch trial. All trials were randomized.

#### 2.3.2 Illusory Double Flash Task

The task was also performed separately for each eye with a randomized order, with the non-tested eye covered by an eyepatch. Participants reported the number of perceived flashes (0–3). The F1B2 condition was presented ten times per stimulus location, and the F1B1 condition served as a single catch trial. All trials were randomized.

#### 2.3.3 Beep Detection Task

Only age-matched controls performed this task, viewing binocularly. Participants reported the number of beeps heard (0–3). All six audiovisual combinations of flashes (0 or 1) and beeps (0, 1, or 2) were tested three times per stimulus location in randomized order.

### 2.4 Experimental setup

Stimuli were presented using Psychtoolbox 3.0.17 in MATLAB 2021a. Auditory tones (62 dB SPL) were delivered via stereo speakers (Bose Companion 2 Series III) placed beside the monitor. Visual stimuli appeared on a 27-inch LCD display (Dell UltraSharp U2720Q; 60 Hz). Participants sat 50 cm from the display with their head stabilized on a chin rest and the non-tested eye occluded. The experiment took place in a darkened room. All experimental scripts are publicly available under a CC-BY license [Chan, 2025].

### 2.5 Data analysis

#### 2.5.1 Average Perceived Flash Count by Task and Stimulus Location

To visualize overall perceptual patterns, we first computed the mean number of flashes reported at each stimulus location for both the Visual Flash Detection (F1B0 trials) and Illusory Double Flash (F1B2 trials) tasks (Fig. 2). Locations were further categorized individually for each participant as visible (flash detection accuracy *>* 50%) or invisible (≤ 50%) based on performance in the detection task. Two Wilcoxon signed-rank tests were conducted within each group (low vision, sighted control) to compare perceived flash counts between tasks (all stimulus locations and both eyes’ data pooled). To control for multiple comparisons, Bonferroni correction was applied across the two tests.

**Figure 2:**
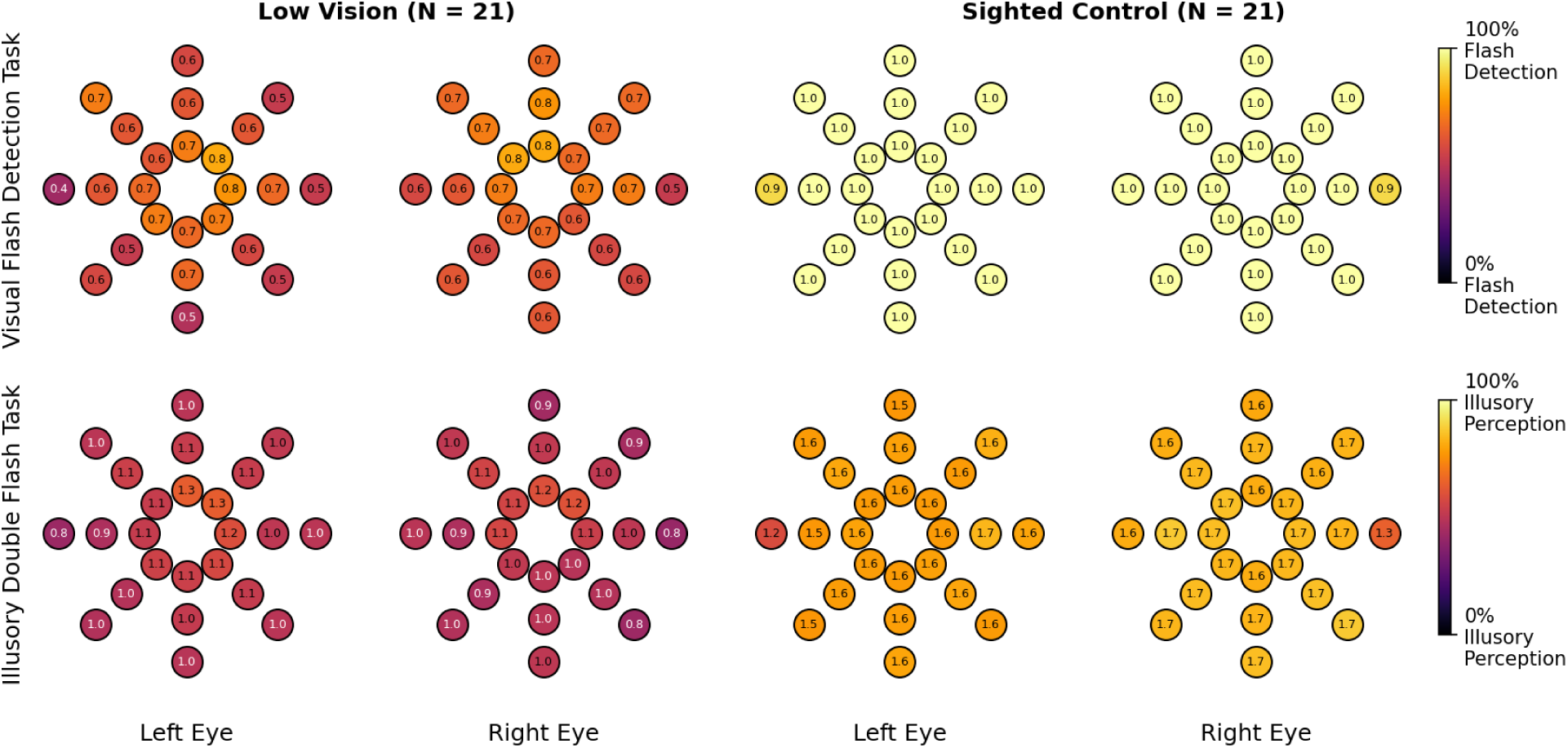
Mean number of flashes perceived by task across all stimulus locations. Top row: *Visual Flash Detection task* (F1B0 condition). Bottom row: *Illusory Double Flash task* (F1B2 condition). Left panel: Low-vision group. Right panel: Sighted control group. Stimulus locations were further categorized for each participant as **visible** (flash detection accuracy *>* 50%) or **invisible** (≤ 50%) based on performance in the detection task.

#### 2.5.2 Accuracy

Accuracy was defined as the proportion of trials in which participants correctly reported the true number of flashes presented (Fig. 3A). To examine how visual and spatial factors influenced visual accuracy, we fit two separate linear mixed-effects models using the statsmodels package in Python. In both models, the dependent variable was mean flash perception accuracy and we included a random intercept for participants to account for individual differences in overall accuracy. The first model tested the effects of *group* (Low Vision, Sighted Control), *condition* (F0B0, F1B0, F1B1, F1B2), and *visibility* (Visible, Invisible), along with all interactions among these factors. The second model tested the effects of *group*, *condition*, and spatial *eccentricity* (5*^◦^,* 10*^◦^,* 15*^◦^*), and their interactions.

**Figure 3:**
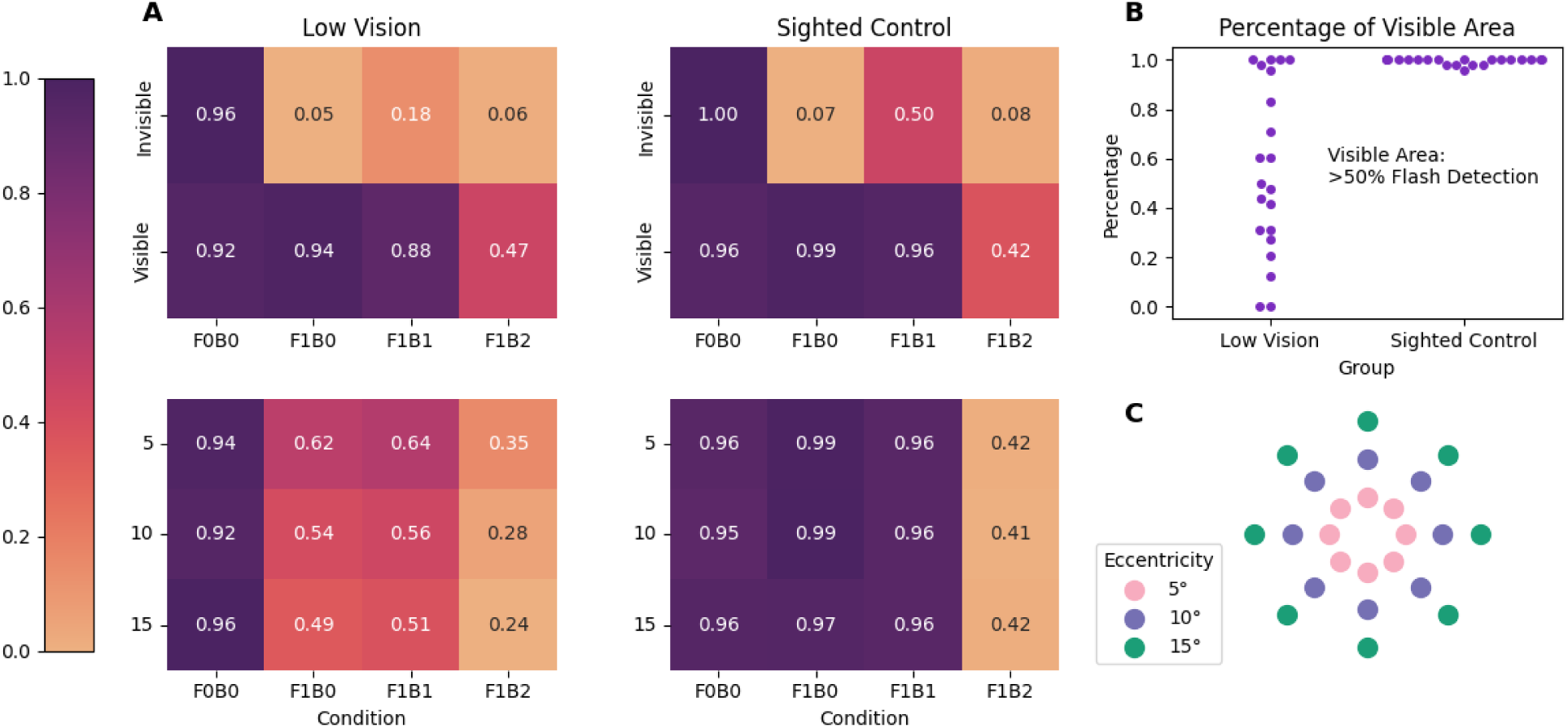
Mean accuracy of flash perception across conditions. **(A)** Heatmaps show accuracy in reporting the true number of flashes (0/1) across conditions and locations classified as *visible* (flash-detection accuracy *>* 50%) or *invisible* (≤ 50%) based on each participant’s performance in the Visual Flash Detection Task (top panels). Bottom panels show accuracy as a function of stimulus eccentricity (5*^◦^*, 10*^◦^*, 15*^◦^*). F0B0 trials yielded the highest accuracy, indicating correct reports of zero flashes. **(B)** Proportion of stimulus locations classified as visible for each participant. **(C)** Spatial layout of stimulus locations in the task, color-coded by eccentricity (5*^◦^* = pink, 10*^◦^* = purple, 15*^◦^* = green), for illustration purposes.

#### 2.5.3 Flash Detection vs. Perceived Flash Count (F1B1, F1B2)

We examined whether sounds influenced perceived flash numerosity correlates with visual sensitivity (i.e., flash detection accuracy in the F1B0 condition). For each participant, eye, and stimulus location, we first computed flash-detection accuracy in the F1B0 condition, yielding 48 data points per participant (24 locations × 2 eyes). We then computed, separately for each beep numerosity (B1, B2), the mean number of flashes perceived at each location and eye, again yielding 48 data points per beep condition. To assess the relation between visual sensitivity and sound-driven changes in perceived flash numerosity, we correlated flash-detection accuracy with the mean number of flashes reported using Pearson’s correlation coefficient, computed separately for each group (Low-Vision, Sighted Control) and beep condition (B1, B2).

#### 2.5.4 Bayesian Causal Inference Model

Participants’ responses were modeled using a Bayesian Causal Inference (BCI) framework [Körding et al., 2007], from which the forced-fusion, full-segregation, and maximum-likelihood estimation (MLE) models [Ernst and Banks, 2002] were derived as special cases.

In the BCI model, each sensory event (*s_i_*) gives rise to a noisy observation (*x_i_*), sampled from a Gaussian centered on the true stimulus value with modality-specific uncertainty (*σ_V_*, *σ_A_*; Eq. 1). Observers also hold a prior expectation about stimulus numerosity, modeled as a Gaussian with mean *µ_P_* and standard deviation *σ_P_* (Eq. 2).

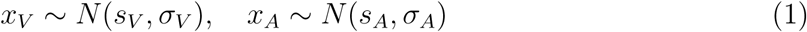

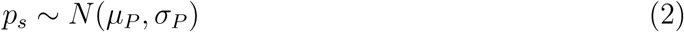

The model infers whether visual and auditory signals arise from a common (*C* = 1) or independent (*C* = 2) cause by combining sensory likelihoods with a causal prior (*p*_common_) via Bayes’ rule (Eq. 3). When signals are inferred to share a common cause, the perceived numerosity is a reliability-weighted average of the visual, auditory, and prior estimates; if inferred as independent, each modality is estimated separately.

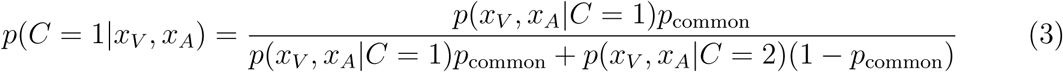

Observers combine these estimates across causal structures (*C* = 1, 2) using one of three decision strategies: *model averaging* (weighted by posterior probability), *model selection* (choosing the more probable cause), or *probability matching* (stochastic selection in proportion to posterior probability) [Körding et al., 2007].

Auditory noise (*σ_A_*) was estimated separately because only the age-matched control group completed the beep-detection task. For each control participant, we fit a reduced BCI model to auditory-only trials, assuming independent causes (*p*_common_ = 0) and using the model-selection strategy (i.e., unisensory estimation). Visual and prior parameters were fixed (*σ_V_* = 0.4, *µ_P_* = 0, *σ_P_* = 4000), leaving *σ_A_* as the only free parameter. Model fits were obtained by maximizing the log-likelihood of observed auditory responses. The mean fitted *σ_A_* across participants was 0.2, which was subsequently fixed in the main model fits.

To further reduce the number of free parameters, the prior mean (*µ_P_*) was fixed at 1.5, based on previous work on multisensory numerosity judgments [Wozny et al., 2008]. We compared four models: (i) the full BCI model, (ii) forced-fusion (*p*_common_ = 1), (iii) full-segregation (*p*_common_ = 0), and (iv) maximum-likelihood estimation (MLE), assuming a common cause with a flat prior (*σ_P_* = 4000, *µ_P_* = 1.5). Free parameters for each model are summarized in Table 2.

**Table 2:** Overview of models and their free parameters.

| Model | Free Parameters | # |
| --- | --- | --- |
| Bayesian Causal Inference (BCI) | $p_{\text{common}}, \sigma_P, \sigma_V$ | 3 |
| Forced-fusion | $\sigma_P, \sigma_V$ | 2 |
| Full-segregation | $\sigma_P, \sigma_V$ | 2 |
| Maximum-likelihood estimation (MLE) | $\sigma_V$ | 1 |

## 3 Results

### 3.1 Average Perceived Flash Count

Participants perceived more flashes during the Illusory Double Flash (F1B2) task (Low Vision: 1.03 ± 0.15; Sighted Control: 1.61 ± 0.1) than during the Visual Flash Detection (F1B0) task (Low Vision: 0.55 ± 0.07; Sighted Control: 0.98 ± 0.01), consistent with an auditory-induced enhancement of visual perception (Fig. 2). This effect was significant in both groups (both Bonferroni-corrected *p <* .001). However, only sighted control participants exhibited the full Double Flash Illusion, reporting an average of 1.61 ± 0.1 flashes across all stimulus locations, whereas low vision participants reported an average of 1.03 ± 0.15 flashes.

Together, these findings indicate that while auditory cues increased perceived flash counts in both groups, the canonical Double Flash Illusion—perceiving two flashes in response to one flash paired with two beeps—was reliably experienced only by sighted controls.

### 3.2 Accuracy

We defined accuracy as the proportion of trials in which participants correctly reported the true number of *flashes* presented (0 or 1; Fig. 3A). To examine how auditory context, spatial location, and group membership influenced accuracy, we fit two linear mixed-effects models with *participant identity* included as a random effect to account for baseline differences in accuracy across individuals. Both models had *accuracy* as the dependent variable, where fixed effects included *group identity* (Low Vision, Sighted Control), *condition* (F0B0, F1B0, F1B1, F1B2), and *stimulus locations* (Model One: visible and invisible locations, see Fig. 3B; Model Two: eccentricity—5*^◦^,* 10*^◦^,* 15*^◦^*, see Fig. 3C).

Accuracy differed systematically across auditory conditions and visibility levels (Fig. 3B). The mixed-effects model revealed a significant main effect of condition (*p <* .001), with participants performing most accurately in the no-flash baseline (F0B0) and showing progressively lower accuracy as the number of accompanying beeps increased (F1B0 *>* F1B1 *>* F1B2). Visible locations yielded slightly higher accuracy than invisible ones, though this main effect of visibility did not reach significance (*p* = .28), likely due to similar levels of accuracy for F0B0 conditions. However, visibility strongly interacted with condition (all *p <* .001): accuracy was significantly higher for visible than invisible locations when one flash is physically present (F1B0: *β* = 0.91, *p <* .001; F1B1: *β* = 0.66, *p <* .001; F1B2: *β* = 0.42, *p <* .001). No significant main effects or interactions involving group were observed, indicating that low vision and sighted control participants showed similar patterns of performance.

We next examined how flash detection accuracy varied with spatial eccentricity (Fig. 3C). The mixed-effects model revealed a significant main effect of condition (*p <* .05), replicating the pattern observed in the visibility analysis: accuracy was highest in the no-flash baseline (F0B0) and declined as the number of beeps increased (F1B2 *<* F1B1 *<* F1B0). In contrast, neither eccentricity nor its interactions with condition or group reached significance (all *p >* .07), indicating that performance remained stable across 5*^◦^*, 10*^◦^*, and 15*^◦^* locations. A modest interaction between group and condition emerged for the F1B0 condition (*β* = 0.30, *p* = .024), suggesting slightly higher accuracy for sighted controls than low vision participants in the baseline task, though this effect was not observed in the other conditions.

Beep-detection accuracy was near ceiling across all audiovisual conditions. Because this task was performed only by a subset of sighted control participants, it was not included in the main analyses. The corresponding heatmap is presented in Supplementary Figure S1.

Together, these results indicate that flash numerosity accuracy was primarily determined by auditory context (highest when there is no sound influence) and visibility of the stimulus location rather than by viewing eccentricity. Both low vision and sighted participants showed reduced accuracy when auditory beeps accompanied flashes, but similar spatial patterns of performance across the visual field.

### 3.3 Flash Detection vs. Perceived Flash Count (F1B1, F1B2)

We next examined whether individual differences in flash detection accuracy correlated with the strength of sound-induced flash perception. For each participant, eye, and stimulus location, flash detection accuracy in the F1B0 condition was correlated with the mean number of flashes reported in the one-beep (F1B1) and two-beep (F1B2) conditions (Fig. 4).

**Figure 4:**
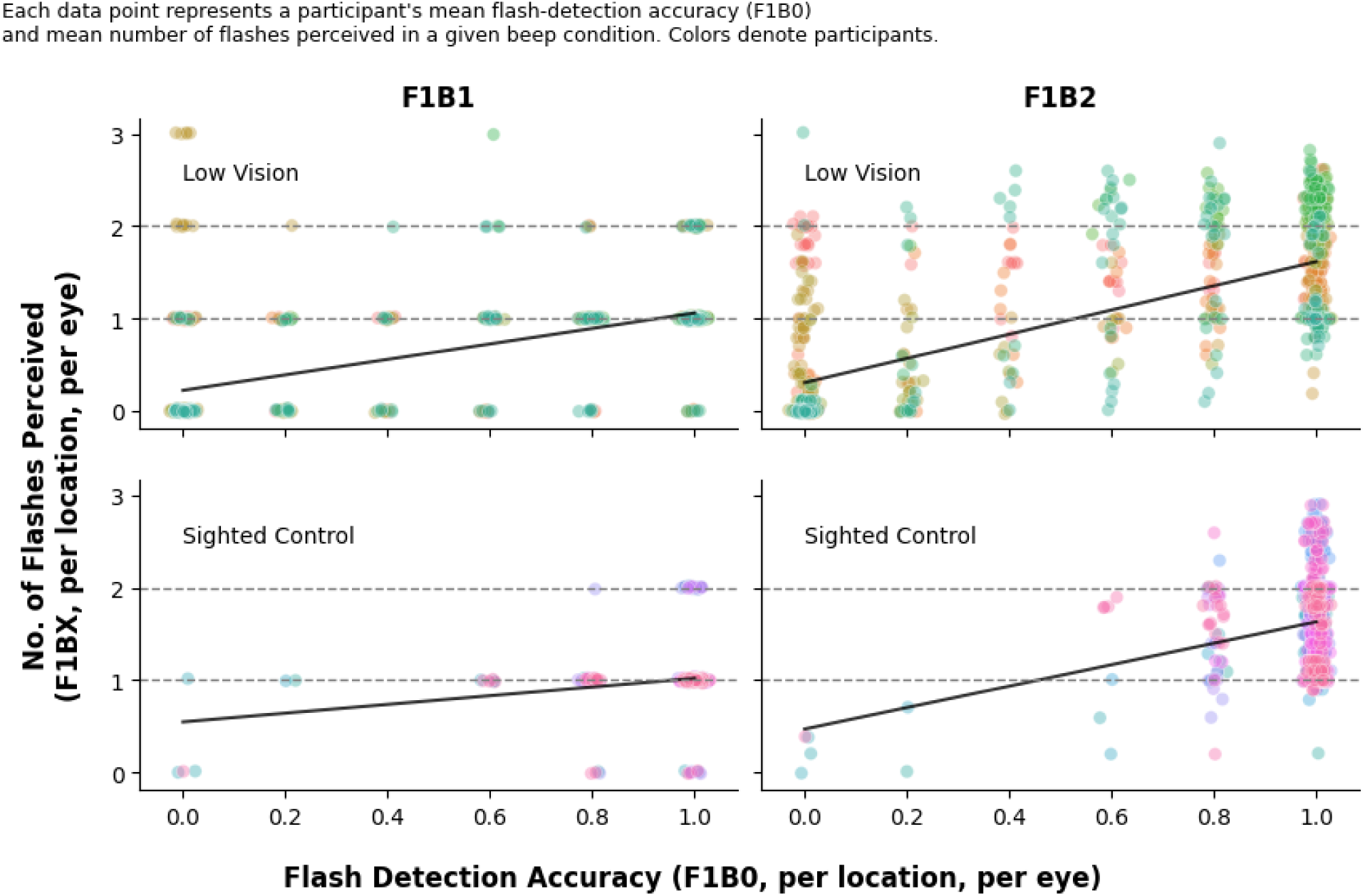
Correlation between flash detection accuracy and perceived flash counts. Scatter plots show the relationship between flash-detection accuracy (F1B0; Visual Flash Detection Task) and the average number of flashes perceived in the one-beep (F1B1, left) and two-beep (F1B2, right) conditions in the Illusory Double Flash Task. Each data point corresponds to a single participant’s mean flash-detection accuracy and perceived flash count at a given stimulus location and eye (48 data points per participant: 24 locations × 2 eyes). Top row: Low Vision group; Bottom row: Sighted Control group.

Both groups showed strong and positive correlations between flash detection accuracy and perceived flash numerosity for both one-beep (Low Vision: *r* = 0.65, *p <* .001; Sighted Control: *r* = 0.21, *p <* .001) and two-beep (Low Vision: *r* = 0.70, *p <* .001; Sighted Control: *r* = 0.21, *p <* .001) conditions. However, visual inspection revealed substantial in-dividual variability, particularly among low vision participants, suggesting that while higher flash detection accuracy generally predicted stronger sound-induced flash enhancement, the strength of this relationship varied markedly across observers.

Together, these results indicate that auditory-driven increases in perceived flash numerosity were present in both groups but expressed with considerable individual variability in magnitude.

### 3.4 Bayesian Causal Inference Model

#### 3.4.1 Model Comparison

We compared model performance across the Bayesian Causal Inference (BCI), forced-fusion, full-segregation, and Maximum Likelihood Estimation (MLE) models using the Bayesian Information Criterion (BIC). For each participant and stimulus location, the model with the lowest BIC value was taken to represent the best trade-off between model fit and complexity. Across participants, the BCI model consistently yielded lower BIC scores than the alternative models, indicating superior explanatory power for both groups (Sighted Control: 238.56 ± 38.18; Low Vision: 965.91 ± 134.38). In contrast, the forced-fusion (Sighted Control: 326.62 ± 45.72; Low Vision: 1129.88 ± 142.47), full-segregation (Sighted Control: 388.38 ± 46.15; Low Vision: 1067.16 ± 135.94), and MLE (Sighted Control: 319.72 ± 45.73; Low Vision: 1122.04 ± 142.33) models provided substantially poorer fits.

Model fits were overall weaker for low vision participants compared to sighted controls, as reflected in the coefficient of determination (*R*^2^). For the BCI model, *R*^2^ values averaged 0.99 for sighted controls and 0.87 ± 0.03 for low vision participants, suggesting greater interparticipant variability and noisier data among the latter group.

If the model’s goodness of fit is poor, the data cannot be said to be adequately explained, and the modeling attempt must be considered unsuccessful [Odegaard et al., 2017, Nagelkerke, 1991]. We therefore adopted a threshold of *R*^2^ ≥ 0.8 to indicate a high-quality fit. For all subsequent analyses, only participants whose model fits met this criterion were included, resulting in 13 Low Vision participants and all participants in the Sighted Control group.

Model performance, as indexed by BIC scores, was strongly and negatively correlated with flash detection accuracy for the Low Vision group (Fig. 5). This relationship was robust both before (*r* = −0.93, *p <* .001) and after excluding participants with poor fits (*r* = −0.93, *p <* .001), indicating that participants with higher flash detection accuracy tended to exhibit better model fits (lower BIC scores). In contrast, the correlation for sighted controls was not significant (*r* = −0.39, *p* = .08). No significant correlation was found between disease duration and age of onset among low vision participants (Supplementary Figure S2).

**Figure 5:**
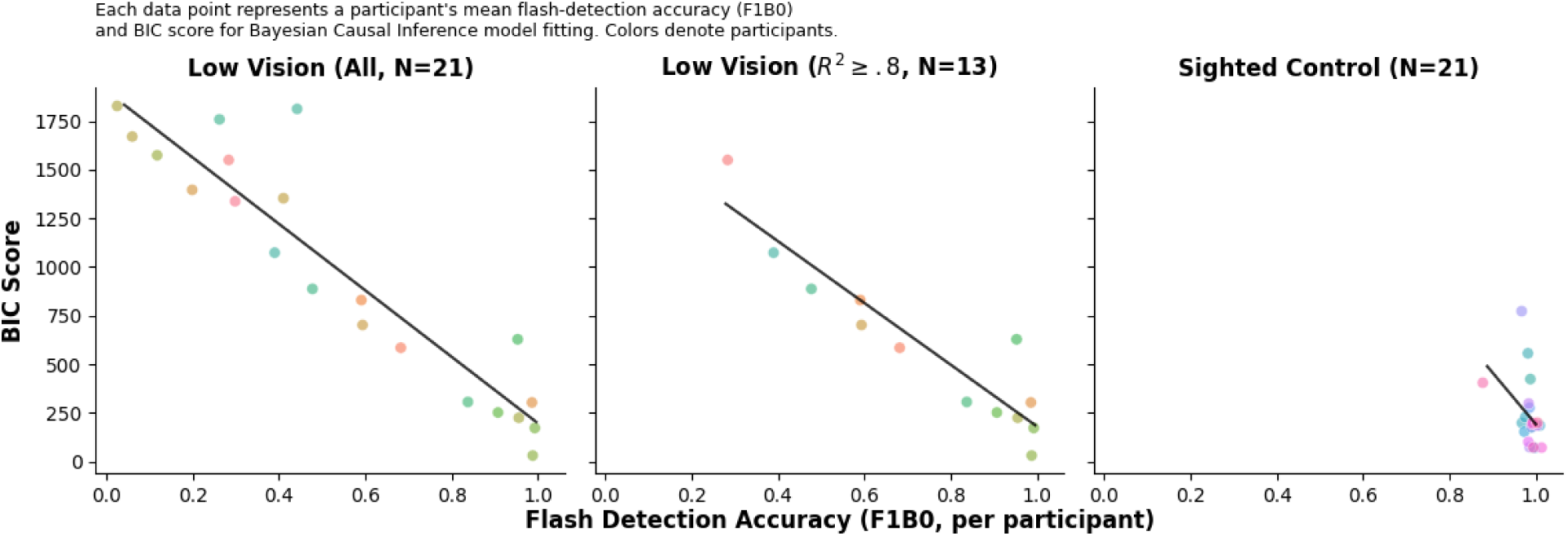
Correlation between flash detection accuracy and BIC scores (Model performance; the lower the better). Each data point shows a participant’s mean flash-detection accuracy (F1B0; all locations pooled) versus their BIC score from Bayesian Causal Inference model fitting. Points are color-coded by participant. Left: All low vision participants (N=21). Middle: Low vision participants with high-quality fits (*R*^2^ ≥ 0.8, N=13). Right: Sighted controls (N=21).

#### 3.4.2 BCI Model Estimated Parameters

We fit a Bayesian Causal Inference (BCI) model to each participant’s data to estimate (1) the prior tendency to assume a common cause (*p_common_*), (2) the precision of the prior over numerosity (*σ_P_*), and (3) the sensory noise of visual flashes (*σ_V_*). Together, these parameters describe how strongly a participant integrates auditory and visual inputs and how reliably they perceive flash numerosity.

Because the BCI framework allows different decision rules for translating posterior distributions into discrete responses, we evaluated three standard strategies: model averaging, model selection, and probability matching, for each participant. The *best-fitting* strategy was chosen based on BIC, allowing us to capture differences in how participants used multisensory evidence.

Across all participants, model averaging was the most common decision rule (59%), indicating that most observers combined the common-cause and independent-cause estimates rather than committing to a single interpretation per trial. Model selection (24%) and probability matching (18%) were less frequently favored.

After excluding participants with poor overall fits (see previous section), the resulting parameter estimates for low-vision and sighted groups are summarized in Table 3. Mann–Whitney U tests revealed no significant group differences in any parameter, suggesting similar inferred prior beliefs (*p_common_*), sensory noise levels (*σ_V_*), and prior variance (*σ_P_*) across groups. Given the high variability and generally weak fits among low vision participants, direct group comparisons should be interpreted with caution. Representative model fits for two example participants are shown in Supplementary Figure S3.

**Table 3:** Parameter estimates (mean ± SE) across participants.

| Group | Parameters |  |  |
| --- | --- | --- | --- |
| | $p_{common}$ | $\sigma_p$ | $\sigma_V$ |
| Sighted Control (N=21) | 0.81 ± 0.06 | 1.64 ± 0.21 | 0.35 ± 0.03 |
| Low Vision (N=13) | 0.63 ± 0.11 | 1.04 ± 0.23 | 0.32 ± 0.05 |

## 4 Discussion

We investigated whether low vision alters susceptibility to the Double Flash Illusion [Shams et al., 2000] and whether illusion strength depends on local variations in visual reliability. We initially hypothesized in line with the “sensory compensation hypothesis” [Braun, 2016], where we predicted stronger illusions because weakened visual input increases reliance on auditory cues. However, it can also be interpreted as multisensory effects should depend on the *availability* (in addition to the *reliability)* of local visual input, and thus may be reduced when visual signals are weak. Our data supports the second interpretation.

Contrary to our initial prediction, low vision participants were not more susceptible to the illusion than sighted controls. Both groups showed the typical auditory-driven increase in perceived flash numerosity, but only sighted controls reliably perceived a full illusory second flash (Fig. 2, 3A). Low vision participants exhibited modest auditory-driven enhancement but did not generate the categorical percept of “two flashes”. Thus, reduced visual function did not amplify audiovisual interactions; instead, illusion strength was gated by whether a sufficiently reliable visual signal was present.

Illusion strength did not vary with eccentricity. Instead, perceived numerosity closely tracked *local flash detection accuracy*. Locations with higher flash detection accuracy were precisely those where participants perceived more flashes when paired with beeps. This supports the interpretation that sound-induced enhancements rely on the presence of a clear visual signal—or at minimum a strong prior belief that a flash occurred—and do not arise when visual input is too weak or inconsistent [Rahnev and Denison, 2018].

Bayesian Causal Inference modeling provided converging but nuanced evidence. While the BCI model outperformed alternative models [Körding et al., 2007, Shams and Beierholm, 2022], fit quality was markedly weaker and more variable in the low vision group (Fig. 5). BIC scores were strongly associated with flash-detection accuracy, indicating that participants with poorer flash detection accuracy deviated more from Bayesian-optimal predictions. This likely reflects a combination of increased internal noise, lapses, inconsistent decision strategies, or reliance on heuristics rather than probabilistic integration [Knill and Pouget, 2004].

A potential concern is that BCI models are explicitly designed to handle a wide range of sensory noise, so why should noise impair model fits? The key issue is not simply *magnitude* of noise, but the *nature* of variability in low vision. The model assumes consistent Gaussian noise, priors, and decision rules across trials. In contrast, low vision observers often exhibit (1) location-specific fluctuations in detectability, (2) intermittent failures to perceive the flash at all, (3) heterogeneous use of strategies, and (4) occasional lapses or guessing patterns that violate model assumptions. Moreover, without precise retinal localization (due to lack of eye-tracking calibrated for low vision), the model receives less informative sensory inputs than intended. These factors reduce the identifiability of parameters and lead to poorer fits even though the model formally allows high-noise solutions.

Despite these differences in fit quality, the estimated parameters showed no significant group differences. In particular, the visual-noise parameter (*σ_V_*) was unexpectedly similar across groups. Given the clear behavioral differences and the reduced fit quality in the low vision group, this similarity likely reflects a limitation of the model rather than genuine equivalence in sensory precision. When fits are poor, parameter estimates become less interpretable due to trade-offs or non-identifiability, and group comparisons should be interpreted with caution.

Finally, our behavioral findings refine, rather than contradict, neural evidence showing cross-modal recruitment of occipital cortex in visual impairment [Cunningham et al., 2015, Collignon et al., 2009, Kupers and Ptito, 2014]. Despite such reorganization, low vision observers did not exhibit globally enhanced multisensory interactions. Instead, auditory influence on perceived numerosity was tightly constrained by the reliability of local visual input.

Together, these results show that degraded vision does not automatically amplify audiovisual interactions. Sound-induced increases in perceived flash count emerge only when visual input is sufficiently reliable, and poorer vision leads to weaker and more variable alignment with Bayesian models. The Double Flash Illusion reflects a flash-detection-accuracy-dependent interaction, and group-level BCI parameter comparisons should be interpreted within the limits of model adequacy and fit quality.

An additional consideration is the heterogeneity of our low vision cohort. Participants differed in diagnosis, spatial extent of field loss, and disease duration, contributing to the variability observed in both behavior and model fits. Although eye tracking would have allowed direct verification of fixation stability and retinal locus of stimulation, standard calibration is often infeasible for low vision individuals. At the time of data collection, our lab did not yet have accessible calibration tools (e.g., skipped-calibration modes, non-foveal target methods). We therefore relied on task structure and instructions rather than direct gaze measurement. Future work incorporating low-vision-adaptive eye tracking will be crucial for linking local retinal function, fixation behavior, and multisensory integration with higher fidelity.

## 5 Conclusion

This study examined how reduced visual function shapes audiovisual interactions in the Double Flash Illusion. Contrary to a simple sensory compensation account, low vision did not lead to enhanced illusion susceptibility. Instead, auditory-driven increases in perceived flash numerosity depended critically on the availability and reliability of local visual input. Sound influenced perception only when a visual signal was sufficiently detectable, and illusion strength closely tracked flash detection accuracy rather than eccentricity or group membership.

Bayesian Causal Inference modeling provided converging support for this interpretation while highlighting important limitations. Although the BCI framework captured overall response patterns, model fits were weaker and more variable in low vision participants, particularly at locations with poor flash detectability. Under these conditions, parameter estimates were less identifiable and group-level comparisons were therefore not informative. These findings underscore that deviations from Bayesian predictions in low vision reflect not merely increased sensory noise, but heterogeneity in perceptual reliability, decision strategies, and model assumption violations.

Together, our results demonstrate that degraded vision does not automatically amplify multisensory integration. Instead, audiovisual interactions are constrained by the presence of a usable visual signal, even in the context of long-term visual impairment and potential neural reorganization. The Double Flash Illusion thus reflects a detection-dependent multisensory process rather than a globally enhanced compensatory mechanism. Future work combining low-vision-adaptive eye tracking with computational models that allow trial-level variability and lapses will be essential for more precisely characterizing multisensory inference under sensory uncertainty.

## Acknowledgements

We are grateful for support from the National Institutes of Health, National Eye Institute (R01EY031761); the National Institutes of Health, the BRAIN Initiative (5K99EY031987-02). A.C. thanks the Croucher Foundation for a scholarship.

## 6 Author Contributions

N.S. conceived the idea. A.C., N.S. and A.T. designed the experiments. A.C., N.S. and C.L. recruited participants. A.C. carried out the experiments, conducted data analyses and wrote the manuscript with input from all authors. S.S. supervised the project.

## 7 Competing Interests Statement

The authors declare no competing interests.

## Notes

### Competing Interest Statement

The authors have declared no competing interest.

