## Supplementary Information for "Did you see the sound? A Bayesian Perspective on Crossmodal Perception in Low Vision"

**Figure S1. Mean Accuracy of Perceiving Beeps (N=16)**

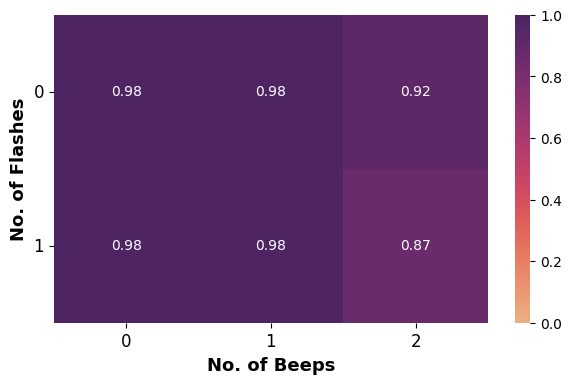

Heatmaps show accuracy in reporting the true number of beeps (0/1/2) across conditions. Performance is near ceiling (1.0) across conditions, suggesting that number of flashes present has little influence on beep detection accuracy.

**Figure S2. Correlation between Disease Characteristics and BIC Scores (Model Performance; the lower the better)**

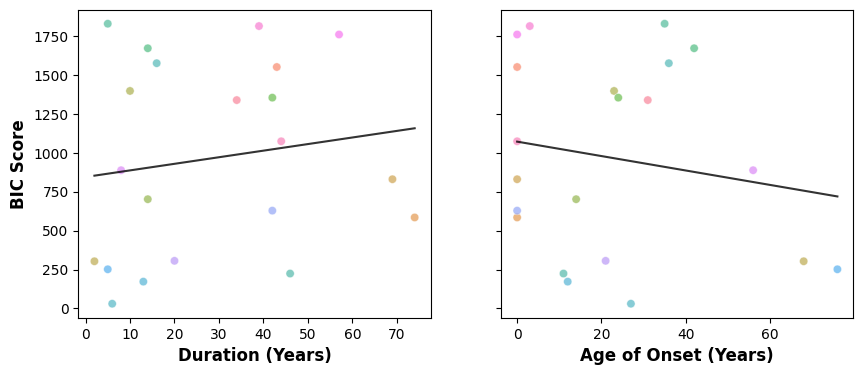

There are no significant correlations between disease duration and BIC score, as well as age of onset and BIC score.

**Figure S3. Example Bayesian Causal Inference model fits for two representative participants.**

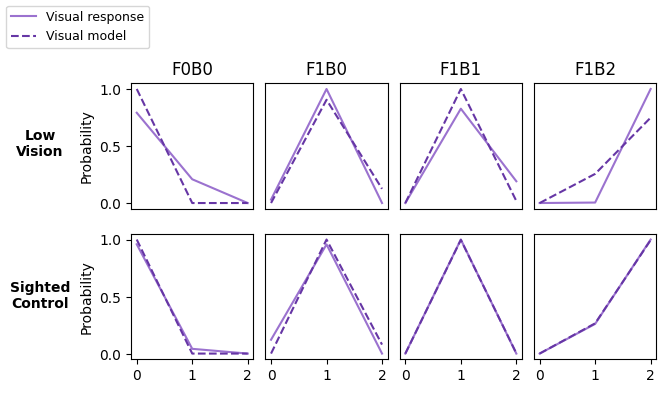

Visual response distributions (solid lines) and corresponding model-predicted distributions (dashed lines) are shown for four conditions (F0B0, F1B0, F1B1, F1B2). The top row depicts a representative participant from the Low Vision group, and the bottom row depicts a Sighted Control participant. Each panel shows the probability of reporting 0–2 flashes for that condition. Overall, the model provides a close approximation to participants’ response profiles, capturing both unimodal visual responses (F0B0, F1B0) and patterns associated with illusory percepts in multisensory conditions (F1B1, F1B2)
